# Machine-learning MRI stratification of genetic frontotemporal dementia for clinical trial enrichment

**DOI:** 10.64898/2026.07.22.740145

**Authors:** Mahdie Soltaninejad, Yasser Iturria-Medina, Arabella Bouzigues, Lucy L. Russell, Phoebe H. Foster, Eve Ferry-Bolder, John C. van Swieten, Lize C. Jiskoot, Harro Seelaar, Raquel Sanchez-Valle, Robert Laforce, Caroline Graff, Daniela Galimberti, Rik Vandenberghe, Alexandre de Mendonça, Giuseppe di Fede, Isabel Santana, Alexander Gerhard, Johannes Levin, Benedetta Nacmias, Markus Otto, Maxime Bertoux, Thibaud Lebouvier, Chris R. Butler, Isabelle Le Ber, Elizabeth Finger, Maria Carmela Tartaglia, Mario Masellis, James B. Rowe, Matthis Synofzik, Fermin Moreno, Barbara Borroni, Jonathan D. Rohrer, Simon Ducharme, GENFI

## Abstract

**BACKGROUND:** Genetic frontotemporal dementia (FTD) shows large differences in symptom profiles, brain atrophy patterns, and progression rate, making clinical trials difficult to design and power. There is a need for biomarkers that can model disease progression, identify biologically distinct groups, and support efficient trial enrichment.

**METHODS:** We applied contrastive trajectory inference (cTI), a machine-learning method, to structural MRI, white matter hyperintensity, and demographic data from 736 participants in the GENFI cohort, including non-carriers and carriers of *C9orf72*, *GRN*, or *MAPT* mutations. cTI produced an individual “genetic FTD progression score” (0–1) and grouped mutation carriers into data-driven subtypes. We tested construct validity using correlations between progression score and cognitive/functional measures, examined subtype differences in brain–behavior coupling, plasma neurofilament light (NfL), and longitudinal decline, and compared cTI-based trial enrichment against age, cortical thickness and NfL using analytic and simulation-based power analyses.

**RESULTS:** Genetic FTD progression scores correlated strongly with global dementia severity and multiple cognitive domains (all p < 0.001), confirming robust clinical scoring. Two mutation-carrier subtypes emerged: a Progressive Track (Subtype 2) with strong associations between progression score and cognitive/functional impairment, rising NfL, and faster longitudinal decline; and a Dissociated Track (Subtype 3) with comparable levels of structural variation but weak or absent clinical and NfL changes, suggesting relative biological stability. Baseline subtype membership added prognostic value for future decline in processing speed and language beyond baseline severity. Notably, for *C9orf72* and *GRN*, cTI-informed enrichment reduced required recruited sample size per arm by about 61–75% compared with unenriched designs, and outperformed enrichment using age, cortical thickness or NfL in both analytic and simulation-based power analyses.

**CONCLUSIONS:** Machine-learning stratification of genetic FTD reveals a progressive and a dissociated disease track and provides individualized progression scores that closely track clinical status. cTI progression scores offer a powerful tool for trial enrichment, enabling smaller, more efficient prevention and early-intervention trials than conventional MRI or NfL markers alone.

## 1 Introduction

Frontotemporal dementia (FTD) is a profoundly heterogeneous neurodegenerative disorder that ranks as a common cause of early-onset dementia in individuals under the age of sixty [1]. The disease presents with a spectrum of prominent behavioral alterations, progressive language deficits, and severe executive function impairments that progressively strip away patient autonomy and create heavy burdens for caregivers [2]. Approximately thirty to fifty percent of FTD cases are highly heritable, primarily driven by autosomal dominant mutations in three major genes: chromosome 9 open reading frame 72 (*C9orf72*), progranulin (*GRN*), and microtubule-associated protein tau *(MAPT*) [2]. These mutations result in the abnormal accumulation of pathological proteins in the brain, including TDP-43 and tau aggregates [2]. A defining characteristic of genetic FTD is its considerable clinical and pathological heterogeneity, which presents as highly variable clinical phenotypes, widely differing progression rates, and diverse patterns of brain atrophy, even among individuals carrying the identical genetic mutation within the same family [3,4]. Despite this vast variability, researchers have established that these varied clinical manifestations are universally preceded by a prolonged asymptomatic phase. During this hidden preclinical stage, insidious biological and structural brain changes silently begin to emerge and accumulate over decades before the onset of overt clinical symptoms [5,6].

Designing and executing effective clinical trials for FTD is severely limited by two major, intertwined obstacles: the underlying biological heterogeneity of the disease and the significant logistical challenges associated with patient recruitment [7–9]. The marked variability in clinical presentation and progression rates complicates the selection of uniform trial endpoints, making it exceptionally difficult to measure treatment effects accurately and reliably across a clinically diverse patient cohort [8,10]. Because FTD is a relatively rare condition compared to Alzheimer’s disease, finding a sufficient number of eligible patients who simultaneously meet strict inclusion criteria and are situated near a trial center is a persistent and limiting bottleneck [8]. Consequently, a substantial percentage of clinical trials face the constant risk of failure due to recruitment difficulties, as only a critically small proportion of the total eligible population actually enters and completes these longitudinal studies [6,11]. The lack of any currently approved therapies to effectively stop or slow disease progression magnifies the urgency of overcoming these multifaceted obstacles. This harsh clinical reality underscores the critical need for innovative, adaptive trial designs that can properly accommodate disease heterogeneity while maximizing the scientific utility of a highly limited patient pool.

Traditional clinical trials that enroll only fully symptomatic but mild FTD patients require unacceptably large sample sizes to detect meaningful therapeutic effects, a problem greatly worsened by the naturally variable rates of cognitive and physical decline among participants [10]. To overcome these severe sample size limitations and improve trial efficiency, clinical trial designs are increasingly turning to targeted enrichment strategies as a viable methodological solution. By proactively incorporating asymptomatic mutation carriers who are demonstrably approaching phenoconversion, researchers can significantly reduce the required sample sizes and shorten the necessary duration of observation [6]. The success of this enrichment approach, however, relies heavily on the availability of robust, reliable biomarkers that can accurately forecast a patient’s proximity to symptom onset and precisely measure active neurodegeneration. Without precise disease advance evaluation, broadly incorporating asymptomatic individuals could unintentionally weaken the treatment signal, as many subjects might not exhibit any measurable clinical decline during the standard trial period. Therefore, leveraging biomarkers to selectively identify individuals at high risk for approaching clinical conversion allows for smaller, vastly more statistically powerful prevention and early symptomatic therapeutic trials [6].

Neuroimaging biomarkers, particularly those derived from structural magnetic resonance imaging (MRI), could potentially play a crucial role in tracking FTD progression and facilitating clinical trial enrichment [10,12,13]. Imaging metrics such as cortical thickness and specific subcortical volumes demonstrate significant, quantifiable structural brain changes occurring up to decades before symptom onset, providing a clear window into early pathology [2,3,5].

Furthermore, recent findings definitively highlight that white matter hyperintensities serve as crucial early markers of neurodegeneration, particularly in *GRN* mutation carriers, often preceding cortical atrophy and even elevations in conventional fluid biomarkers like neurofilament light chain [14]. Specific longitudinal studies have documented the capability of these regional MRI markers, including volumetric changes in the thalamus and frontotemporal networks, to predict clinical conversion and map longitudinal decline in asymptomatic populations [13,15,16]. However, the complex spatial and temporal heterogeneity of these structural brain changes across different genetic mutations demands highly sophisticated analytical approaches rather than generalized models [4,17,18]. Advanced machine learning methods, such as contrastive trajectory inference (cTI), offer a powerful, data-driven solution by extracting multidimensional temporal patterns to estimate disease progression and classify subtypes accurately, cleanly overcoming the limitations of traditional rule-based frameworks [19,20].

In this study, we applied cTI to structural MRI data from the Genetic Frontotemporal Dementia Initiative (GENFI) cohort to develop a personalized disease progression framework for genetic FTD. Our objectives were to: (1) identify and characterize distinct disease subtypes within genetic FTD mutation carriers; (2) generate individualized disease scores that quantify disease severity and proximity to clinical conversion; and (3) evaluate whether cTI-derived disease scores improve clinical trial enrichment efficiency compared to conventional biomarkers including cortical thickness and neurofilament light chain (NfL). We hypothesized that by capturing disease heterogeneity through subtype identification and precise individual progression, cTI-based enrichment would substantially reduce sample size requirements for adequately powered clinical trials, thereby addressing a fundamental barrier to FTD therapeutic development.

## 2 Method

### 2.1 Participants and Data

Participants were recruited from the Genetic Frontotemporal Dementia Initiative (GENFI), a multicenter study of individuals with pathogenic FTD mutations and their first-degree relatives [5]. In the present work, we specifically focused on carriers of *GRN, C9orf72,* or *MAPT* mutations and their non-carrier family members. We included (i) symptomatic mutation carriers meeting established clinical criteria for FTD syndromes, (ii) asymptomatic carriers without clinical manifestations at the time of assessment, and (iii) non-carrier relatives who served as controls.

T1-weighted and T2-weighted MRI scans were acquired on 3T scanners (Siemens Trio/Skyra/Prisma, Philips, and General Electric) across GENFI sites using harmonized structural imaging protocols described in detail elsewhere for this cohort [5,14]. Briefly, volumetric T1-weighted images were obtained with sagittal 3D MPRAGE or equivalent sequences, and T2-weighted images with sagittal 3D fast spin echo sequences, using 1 mm isotropic voxels and whole-brain coverage. NfL was measured in blood samples using single molecule array (Simoa) technology according to standardized GENFI laboratory protocols, and values were used as a fluid biomarker of neurodegeneration [21].

### 2.2 Image Preprocessing

#### 2.2.1 Cortical and subcortical measures

T1-weighted images were processed with FreeSurfer (version 7.1.1) for cortical reconstruction and subcortical segmentation, yielding regional cortical thickness and volumetric measures. Cortical parcellation was performed using the Desikan–Killiany atlas, and subcortical labels were obtained from the automated ASEG module. All FreeSurfer-derived volumetric measures were normalized by each participant’s intracranial volume to account for head-size differences. Further details on this pipeline and quality control procedures are provided in our previous work [13].

#### 2.2.2 White matter hyperintensities

White matter hyperintensities (WMH) were segmented using the BISON framework [22] applied to combined T1- and T2-weighted data, following the preprocessing and supervised segmentation protocol described in our earlier publications [14]. This procedure included standard image correction and normalization steps and produced WMH lesion maps in native space that were subsequently registered to the ICBM152 template. Lobar WMH volumes were then derived using the Hammers atlas in standard space. These measures are defined in a common stereotaxic space, so no additional normalization for intracranial volume was needed.

### 2.3 Data Harmonization using ComBat

To reduce scanner-related variability in our imaging dataset, we applied ComBat, an empirical Bayes harmonization method widely used to remove non-biological batch effects while preserving meaningful biological differences [23]. Scanner differences can introduce artificial shifts in mean and variance across imaging features, which can obscure true disease-related effects and reduce statistical power. ComBat models and removes these scanner effects while adjusting for covariates that should not be altered. In our analysis, age, genetic group, genetic status, sex, and education were included as preserved covariates to ensure that harmonization removed only scanner-specific variability and retained true biological signal.

### 2.4 Data Cleaning and Feature Preparation

Healthy controls were defined within the non-carrier group using a multi-step procedure applied only to non-carrier participants. First, all non-carriers with any clinical impairment (FTLD-CDR-SOB > 0) were removed, and outlier detection for plasma NfL and total WMH volume was then performed on the remaining non-carrier sample. Biomarker values were log-transformed, upper outliers were identified using an interquartile-range–based threshold on the log scale, and non-carriers exceeding these thresholds were excluded, yielding the final healthy control group used for downstream analyses (see Supplementary Methods for full details).

Because the number of visits varied across individuals, a single visit per person was retained for both healthy controls and mutation carriers, selected according to a predefined hierarchy that prioritized availability of clinical ratings and plasma NfL, and then the most recent eligible visit. To account for demographic effects, all imaging and biomarker features were residualized for age, sex, and education using linear regression models fitted in the healthy control group, and the resulting residuals were used as covariate-adjusted measures. Left and right hemisphere features were then merged to reduce dimensionality and improve model stability: cortical thickness measures were averaged across hemispheres, and volumetric measures (subcortical volumes, and regional WMH burden) were summed, while two asymmetry indices (left-to-right WMH and left-to-right cortical thickness ratios) were additionally computed; further details and exact feature counts are provided in the Supplementary Methods.

### 2.5 cTI Modeling

To characterize individual genetic FTD progression and identify putative disease subtypes, we utilized the contrastive trajectory inference (cTI) algorithm implemented within the open-access NeuroPM-box software toolkit [20]. cTI is a machine learning method that extracts pseudotemporal disease trajectories from high-dimensional, cross-sectional data [19]. The algorithm utilizes contrastive principal component analysis (cPCA) to uncover latent patterns enriched in a target population by controlling for normal biological variations and noise present in a background population [24]. In our analysis, we defined 266 non-carriers as the background control group and 140 symptomatic carriers as the target group, performing cPCA using all available features without preselection. Because the input features had already been adjusted for confounding factors in preceding steps, no additional covariable correction, outlier correction, or data imputation were applied within the cTI module. To establish individualized disease progression, the algorithm calculates a Euclidean distance matrix among all subjects in the optimized cPCA space and constructs a minimum spanning tree (MST) to determine the shortest path from each individual to the background population. Based on this path, each subject is assigned an individualized disease progression score, or pseudotime, scaled between 0 and 1, where low scores indicate proximity to the healthy background state’s centroid and high scores indicate an advanced disease state. Concurrently, to uncover disease heterogeneity, cTI applies spectral clustering over the cPC-based distance matrix to group subjects into distinct (and potentially overlapping) subtrajectories that represent different variants of disease progression [19,20].

### 2.6 Evaluation of Disease Progression Model Performance

To evaluate the performance of our neuroimaging-based progression model, we computed Pearson correlations between the cTI-derived disease scores and several standardized neuropsychological and clinical measures in mutation carriers (i.e., non-background participants). These included the Mini-Mental State Examination (MMSE; global cognition), FTLD-CDR Sum of Boxes (global dementia severity in FTD), Trail Making Test–Part B (TMT-B; executive function, task-switching, and cognitive flexibility), Digit Symbol Substitution Test (processing speed, attention, and working memory), Boston Naming Test (language and word retrieval), Verbal Fluency tasks (phonemic and semantic fluency), and the MiniSEA (Mini Social and Emotional Assessment; social and emotional processing) [25,26].

### 2.7 Subtype Characterization and Biological Validation

To evaluate the clinical and biological relevance of the subtypes identified by cTI, we performed a multi-modal validation across four domains: cross-sectional clinical-cognitive profiles, longitudinal progression, and biochemical markers of axonal injury.

#### 2.7.1 Genetic and Clinical Composition

We characterized the distribution of mutation types (*C9orf72, GRN, MAPT*) and clinical diagnoses across groups to identify any phenotypic enrichment.

#### 2.7.2 Clinical and Cognitive Associations

Using Pearson correlations, we assessed the relationship between the neuroimaging-derived disease scores and comprehensive battery covering global cognition (MMSE), functional impairment (FTLD-CDR-SB), language (Boston Naming Test), processing speed (Digit Symbol), executive function (Verbal Fluency, Trail Making Test B), and social cognition (Mini-SEA). These analyses were performed within each mutation-carrier subtype to determine the degree of "brain-behavior coupling", specifically, whether imaging-derived structural changes translated into functional impairment.

#### 2.7.3 Biochemical Validation

To investigate whether the imaging-derived subtypes reflected active neurodegeneration, we compared NfL levels. Recognizing that subtypes may occupy different ranges of the disease spectrum, we planned a severity-matched analysis. Specifically, where subtypes showed overlapping disease scores, we compared NfL levels to determine if subjects with similar levels of structural deviation exhibited different levels of active axonal injury.

#### 2.7.4 Longitudinal Stability and Prognosis

Annual rates of change for clinical and cognitive measures were calculated for participants with available follow-up data. Group differences in progression rates were assessed using Mann-Whitney U tests. Finally, we employed linear regression to test if baseline subtype membership provided incremental predictive value (*ΔR*^2^) for longitudinal decline beyond baseline clinical severity.

### 2.8 Evaluation of Trial Enrichment Efficiency

We evaluated whether cTI-derived disease progression scores can improve clinical trial enrichment compared with conventional biomarkers (cortical thickness and NfL) in genetic FTD. We modeled disease-modifying trials with a two-arm design (active vs placebo), using 2-year change in FTLD-CDR-SOB as the primary endpoint, a two-sided α = 0.05, 80% power, and a targeted 50% reduction in clinical progression. Analyses were performed separately for *GRN*, *MAPT*, and *C9orf72* mutation carriers under a hybrid prevention/early-intervention design in which all symptomatic carriers were eligible, while asymptomatic carriers were included only if they met prespecified enrichment criteria based on baseline biomarker abnormality. For each mutation group, we considered both a baseline cohort including all mutation carriers (symptomatic and asymptomatic) and an enriched hybrid cohort composed of all symptomatic carriers plus asymptomatic carriers meeting the enrichment criterion.

Enrichment was evaluated using four baseline strategies: (i) cTI disease progression score; (ii) cortical thickness (inverted so that higher values indicate greater atrophy); (iii) log-transformed plasma NfL; and (iv) age. For each of three first metrics, an abnormality threshold was defined in non-carrier controls only: we computed the 80th percentile at baseline, and mutation carriers were considered “enriched” if their baseline value was greater than or equal to this control-derived threshold. For age, we considered 45 years as an enrichment threshold. This value represents a mid-life time point preceding the typical mean age of clinical onset in genetic FTD and falls within the range where asymptomatic biomarker and neuroanatomical changes have been reported in some carriers.

For power estimation, we constructed a longitudinal dataset with one baseline and one follow-up per participant. Baseline was defined as the earliest visit with a valid cTI disease score. Follow-up was selected as the visit closest to 24 months after baseline among visits with available FTLD-CDR-SOB, accepting any visit within a ±6-month tolerance window (i.e., 18-30 months after baseline); if no visit fell within this window, the nearest available visit was used. Follow-up time was calculated as the age difference (years) between visits, and 2-year clinical progression was estimated as the observed change in FTLD-CDR-SOB divided by follow-up time, multiplied by two.

As a primary, transparent benchmark, we obtained analytic sample-size estimates treating the 2-year FTLD-CDR-SOB change as the primary endpoint. For each mutation group and enrichment strategy, we estimated the mean 2-year change μ and standard deviation σ in the corresponding recruited cohort. A 50% treatment effect was modeled as an absolute difference *Δ* = 0.50 × *μ*, yielding Cohen’s *d* = *Δ*/ *σ*. The analyzable per-arm sample size was approximated as

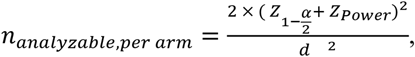

and was then inflated for dropout according to

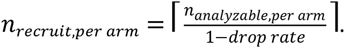

To support and refine these analytic estimates under more realistic assumptions about variance and dropout, we additionally performed Monte Carlo simulation–based power analyses, which are described in detail in the Supplementary Methods.

## 3 Result

### 3.1 Participant Overview

We included 266 non-carriers and 470 mutation carriers across the three genetic groups (*C9orf72, GRN, MAPT*), divided into asymptomatic and symptomatic subgroups. Age, sex, and education were broadly comparable between non-carriers and asymptomatic carriers. Symptomatic carriers were generally older and had lower educational attainment, particularly in the *GRN* and *C9orf72* groups. Adjusted biomarker values (residuals from age-, sex-, and education-matched controls) showed higher NfL, greater white-matter hyperintensity burden and more cortical thinning in symptomatic carriers, consistent with more advanced disease.

**Table 1.** Participant demographics and adjusted biomarker values. The table summarizes demographic characteristics and adjusted biomarker measures for non-carriers and asymptomatic and symptomatic carriers in each genetic group. Adjusted NfL, adjusted WMH, and adjusted mean cortical thickness (left and right hemispheres) are residual values after regressing out age, sex, and education based on the non-carrier control group.

| Group | Non-carriers | C9orf72 |  | GRN |  | MAPT |  |
| --- | --- | --- | --- | --- | --- | --- | --- |
|  |  | Asymptomatic | Symptomatic | Asymptomatic | Symptomatic | Asymptomatic | Symptomatic |
| N | 266 | 135 | 71 | 138 | 42 | 57 | 27 |
| Age (mean ± std) | 46.6 ± 12.9 | 44.5 ± 11.5 | 64.2 ± 7.5 | 47.0 ± 12.3 | 63.9 ± 8.4 | 40.8 ± 11.5 | 57.8 ± 9.1 |
| Sex(M/F) | 112/154 | 55/80 | 43/28 | 50/88 | 18/24 | 23/34 | 17/10 |
| Education(mean ± std) | 14.5 ± 3.3 | 14.3 ± 3.0 | 12.9 ± 3.6 | 14.8 ± 3.4 | 11.9 ± 3.6 | 14.1 ± 3.4 | 13.6 ± 3.6 |
| Adjusted NfL(mean ± std) | 0.0 ± 2.7 | 6.1 ± 19.8 | 33.6 ± 31.1 | 1.8 ± 6.8 | 55.5 ± 38.0 | 1.4 ± 4.7 | 16.3 ± 14.8 |
| Adjusted WMH(mean ± std) | 0.000 ± 7112.877 | 2293.349 ± 11304.997 | 369.851 ± 13103.666 | 983.662 ± 10449.817 | 6908.932 ± 17687.296 | 5407.527 ± 27626.227 | 1745.866 ± 15679.671 |
| Adjusted Mean cortical thickness right(mean ± std) | -0.000 ± 0.076 | -0.061 ± 0.078 | -0.179 ± 0.123 | -0.012 ± 0.071 | -0.151 ± 0.198 | -0.040 ± 0.082 | -0.157 ± 0.097 |
| Adjusted Mean cortical thickness left(mean ± std) | 0.000 ± 0.077 | -0.058 ± 0.080 | -0.197 ± 0.138 | -0.008 ± 0.069 | -0.214 ± 0.165 | -0.037 ± 0.086 | -0.148 ± 0.121 |

### 3.2 Evaluation of Disease Progression Model Performance

Figure 1 summarizes the associations between cTI-derived disease scores and clinical-cognitive measures used to evaluate the model’s performance. Background participants were excluded from the analysis. Each subplot shows a scatter plot with a fitted regression line and the corresponding Pearson correlation coefficient. All correlations were statistically significant (p < 0.001), indicating that higher disease scores were consistently associated with greater global dementia severity (FTLD-CDR-SB) and poorer performance across cognitive and functional domains.

**Figure 1.**
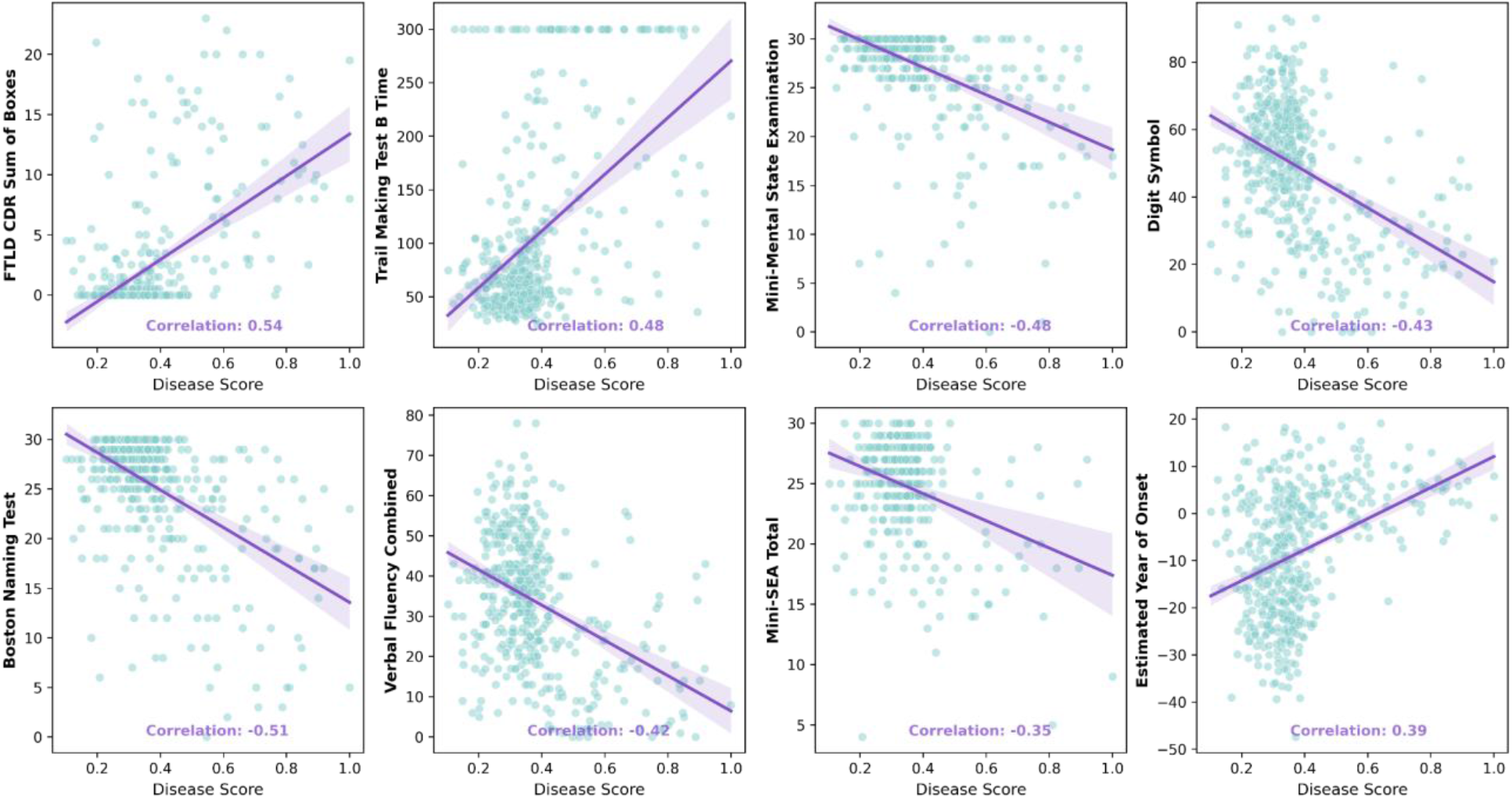
Scatter plot matrix showing the validation of the cTI progression model based on neuropsychological performance. Each subplot presents individual data points along with a fitted regression line and Pearson correlation coefficient (all p < 0.001). Strong negative correlations indicate that higher cTI-derived disease scores correspond to more severe cognitive impairment, confirming the model’s reliability in capturing clinical progression. The shaded areas around the regression lines represent 95% confidence intervals.

### 3.3 Comparative Analysis of Identified Subtypes

The cTI analysis identified three distinct clusters: Subtype 1 (S1), which corresponded to healthy non-carriers, and two mutation-carrier groups, Subtype 2 (S2) and Subtype 3 (S3).

#### 3.3.1 Subtype Distribution and cTI Score Spectrum

We first characterized the distribution of these groups across the disease spectrum (Figure 2). S2 participants spanned the full range of cTI score (0.2 to ∼1.0). In contrast, S3 was confined to the lower cTI score range (0.2–0.5), with a distribution pattern closely resembling the healthy controls (S1). Note that some non-carriers also have non-zero cTI progression scores within the lower part of the scale. This reflects normal variation in brain structure and aging along the latent progression axis, and does not imply clinical or preclinical FTD in healthy individuals.

**Figure 2.**
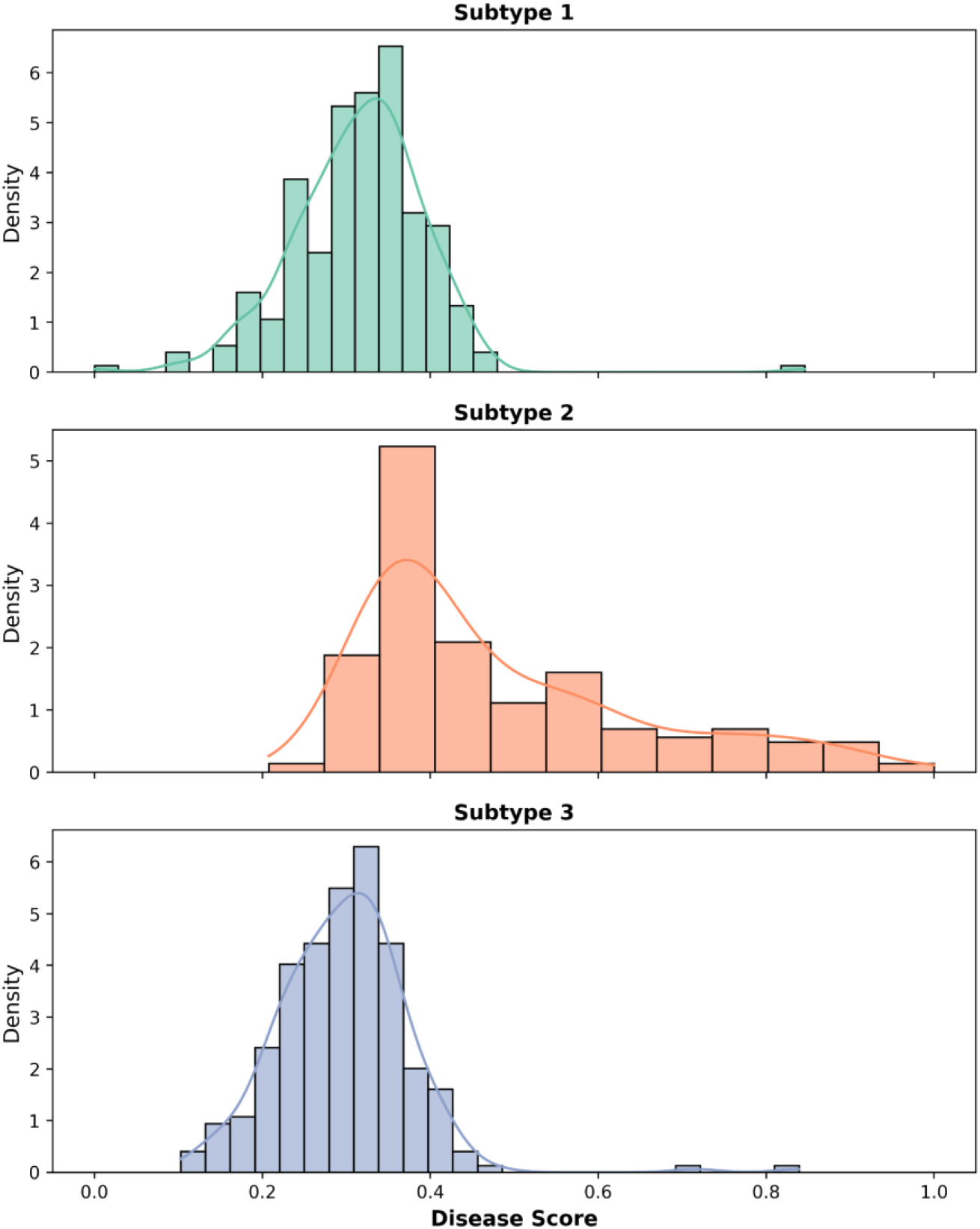
Distribution of cTI progression scores across data-driven subtypes. Histograms show the distribution of cTI progression scores (0–1) in the three subtypes obtained from neuroimaging and demographic data. Subtype 1 (S1) corresponds mainly to healthy non-carriers, whereas subtype 2 (S2) and subtype 3 (S3) consist of mutation carriers. S2 spans almost the full range of cTI scores, while S3 is restricted to lower scores with a distribution similar to S1, indicating profiles closer to the healthy range.

#### 3.3.2 Genetic and Clinical Composition

Both mutation-carrier subtypes included individuals from all three major genetic groups (*C9orf72*, *GRN*, and *MAPT*), but their clinical profiles differed. Subtype 2 had a balanced composition (49.77% asymptomatic and 50.23% symptomatic) and was the primary cluster for symptomatic diagnoses such as bvFTD and PPA, with behavioral variant FTD as the most frequent diagnosis (bvFTD, n=79) followed by smaller numbers of PPA (n=17) and other FTD-related syndromes. In contrast, Subtype 3 was predominantly asymptomatic (87.75%), resulting in much smaller absolute numbers of diagnosed cases, although bvFTD remained the most common diagnosis among its symptomatic minority (n=14). Detailed counts of mutation types and clinical diagnoses per subtype are provided in Figure 3.

**Figure 3.**
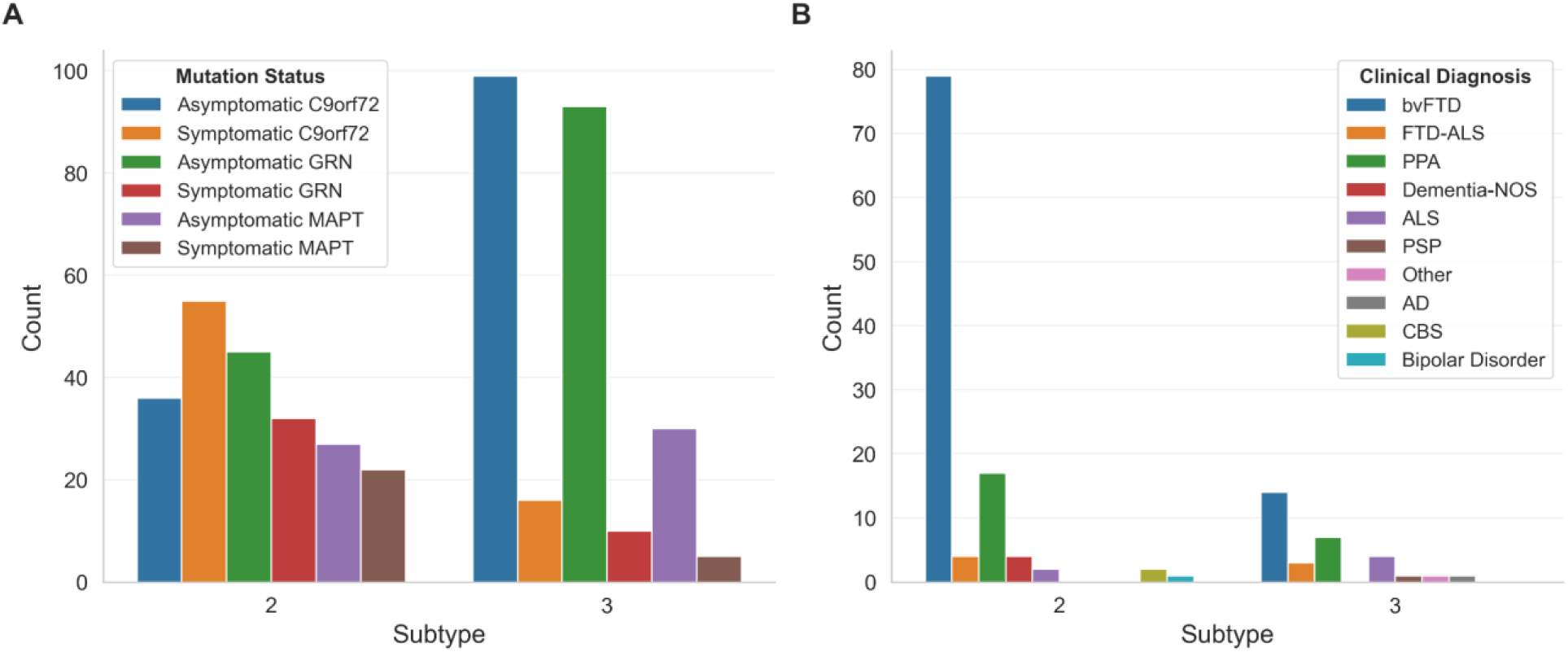
Distribution of genetic mutations and clinical diagnoses across data-driven subtypes. Left panel: Histogram showing, for each subtype, the number of individuals in each genetic group, separated into asymptomatic and symptomatic mutation carriers. Right panel: Histogram showing, for each subtype, the number of symptomatic mutation carriers in each clinical diagnosis category, illustrating how genetic background and presenting syndrome differ between subtypes.

#### 3.3.3 Cross-Domain Clinico-Biological Dissociation

A core finding was the fundamental difference in how neuroanatomical changes translate into clinical symptoms across subtypes (Figure 4).

- **Subtype 2 (The Progressive Track):** Demonstrated robust "brain-behavior coupling" across all domains. Higher disease scores correlated significantly with worse performance in global cognition (MMSE: r=−0.45,p<0.001), functional impairment (CDR-SB: r=0.53,p<0.001), language (Boston Naming Test: r=−0.52,p<0.001), processing speed (Digit Symbol: r=−0.44,p<0.001), executive function (Trail Making Test B time: r=0.45,p<0.001; Verbal Fluency: r=−0.45,p<0.001), and social cognition (Mini-SEA: r=−0.5,p<0.001).
- **Subtype 3 (The Dissociated Track):** Exhibited a striking, across-the-board dissociation. Despite imaging-derived disease scores up to 0.5, no significant correlations were observed for the majority of measures, including Boston Naming Test (r=−0.11,p=0.097), Digit Symbol (r=−0.10,p=0.109), Verbal Fluency (r=−0.02,p=0.813), Mini-SEA (r=0.13,p=0.057), and CDR-SB (r=0.12,p=0.066). This suggests that in S3, the detected structural variations do not translate into clinical symptoms. Only MMSE and Trail Making Test B time showed weak, though statistically significant, correlations (r=−0.16, p=0.014 and r=0.14, p=0.025 respectively), which likely reflect the broader age-related changes observed in this group.

**Figure 4.**
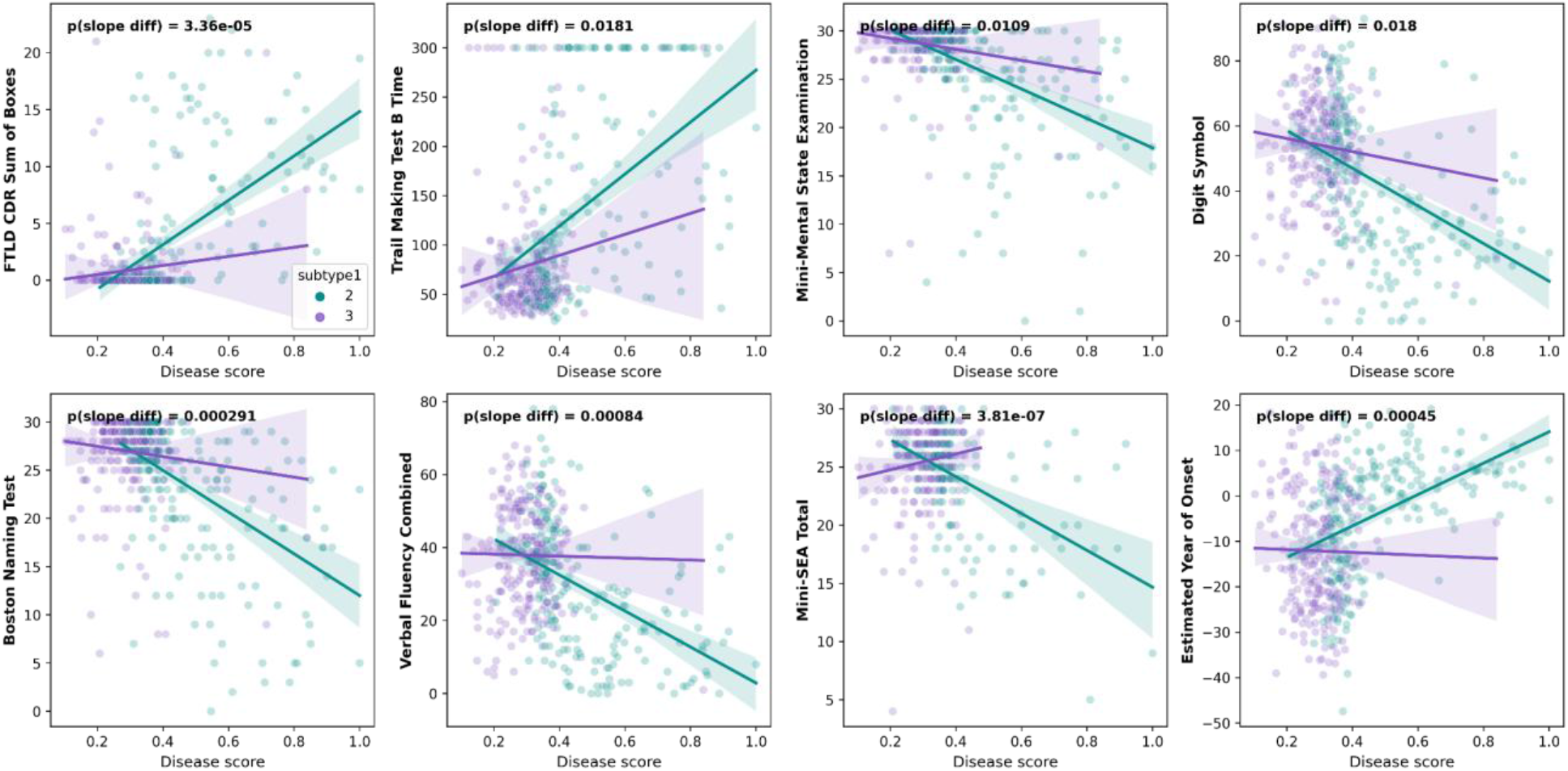
Differential brain–behavior coupling across cTI scores in the Progressive and Dissociated Tracks. Panels show the relationship between cTI score and selected neuropsychological and clinical measures, with subtype 2 (The Progressive Track, green) and subtype 3 (The Dissociated Track, purple) overlaid. For each measure, separate regression lines are fitted for S2 and S3, and the P value for the difference between slopes is reported in each panel; across all measures, slopes in S2 are significantly steeper than in S3, consistent with strong brain–behavior coupling in the Progressive Track and marked dissociation in the Dissociated Track. Shaded regions around the regression lines represent group-specific 95% confidence intervals.

Consistent with this pattern, regression slopes linking cTI progression scores to clinical measures were larger in subtype 2 than in subtype 3 across all domains, indicating stronger brain–behavior coupling in the Progressive Track and relative dissociation in the Dissociated Track (Figure 4).

#### 3.3.4 Severity-Matched Biological Validation (NfL)

Because subtype 3 was restricted to cTI progression scores below 0.5, we compared plasma NfL only among participants from both subtypes within this lower range (cTI progression score < 0.5; Figure 5). Subtype 2 (Progressive Track) participants showed elevated and increasing NfL levels across this range, consistent with ongoing neurodegeneration. In contrast, subtype 3 (Dissociated Track) participants had low, relatively stable NfL levels (around 0–10 pg/mL), and no clear association between cTI progression score and NfL, indicating that their structural brain changes are not accompanied by active axonal injury.

**Figure 5.**
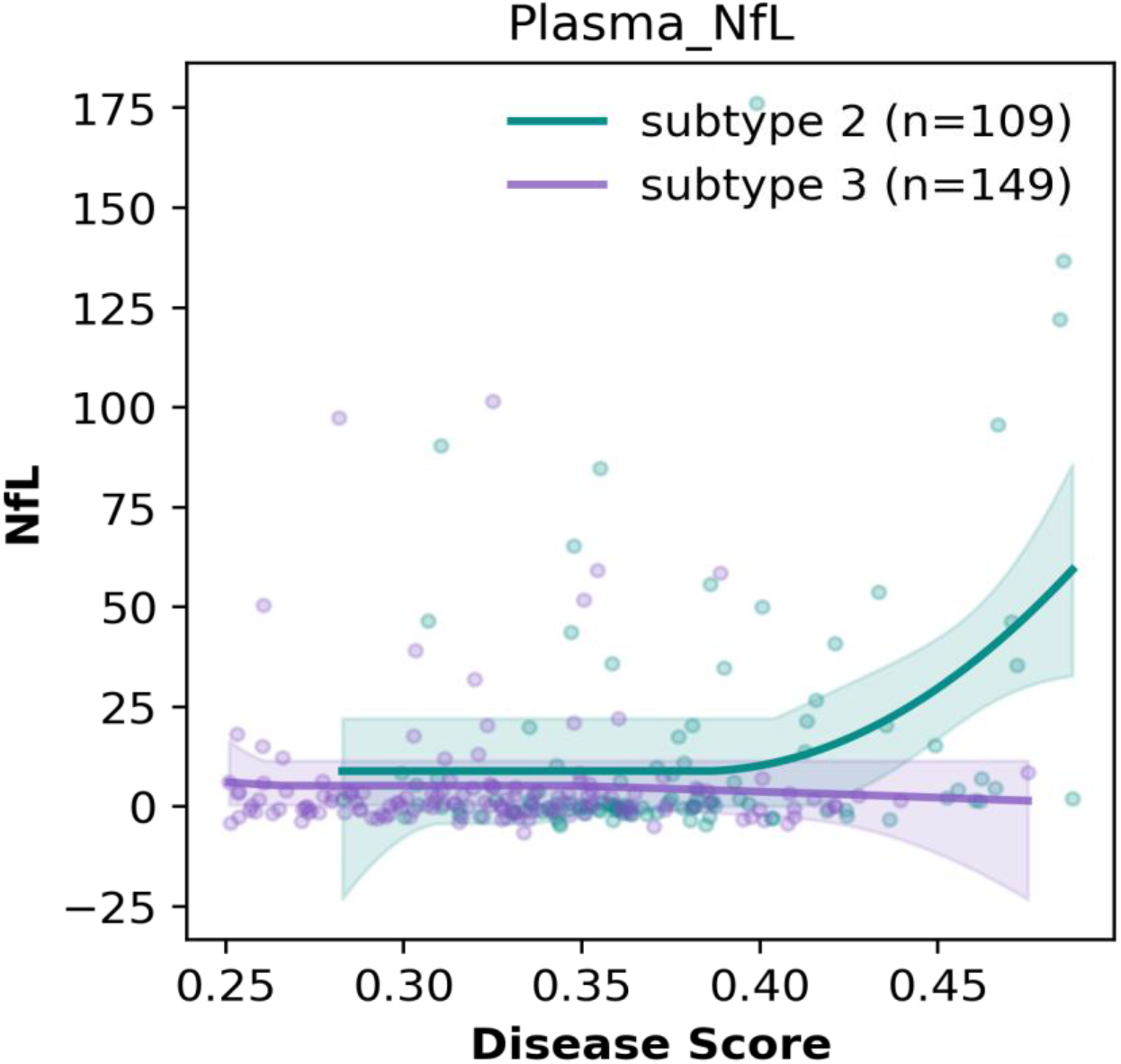
Severity-matched comparison of plasma NfL between Progressive and Dissociated Tracks. Within the shared lower range of cTI progression scores (< 0.5), plasma NfL is plotted for subtype 2 (Progressive Track, green) and subtype 3 (Dissociated Track, purple), with fitted curves and confidence bands for each subtype. Subtype 2 shows increasing NfL with higher cTI progression scores, whereas subtype 3 shows flat, low NfL levels across the same range, supporting active neurodegeneration in the Progressive Track but not in the Dissociated Track.

#### 3.3.5 Longitudinal Progression and Prognostic Value

Longitudinal follow-up confirmed that S2 is an active "Progressive Track," showing significantly higher annual rates of change in MMSE (p=0.0017), CDR-SB (p<0.001), and Digit Symbol performance (p <0.001) compared to S3. The distribution of rates of change for Subtype 2 showed wide variability with substantial departures from zero, indicating active clinical progression. In contrast, S3 participants remained stable, with rates of change tightly centered around zero. Furthermore, baseline subtype membership significantly predicted future decline in processing speed (Digit Symbol: *ΔR*^2^=0.025,p=0.004) and language (*ΔR*^2^=0.013,p=0.035). More details can be found in Supplementary Materials.

### 3.4 Power Analysis

Enrichment thresholds were defined using non-carrier controls, by taking the 80th percentile of each biomarker at baseline (cTI disease score > 0.371, cortical thickness < 2.366 mm, log NfL > 2.417). In the hybrid design, all symptomatic mutation carriers were always eligible, and only asymptomatic carriers who crossed these thresholds were included. This approach created enriched cohorts that were smaller than the full mutation-carrier samples but had more advanced biomarker profiles at baseline.

Analytic power calculations showed that all enrichment strategies reduced the required recruited sample size per arm compared with recruiting all mutation carriers. For *C9orf72*, the unenriched design required 594 participants per arm, whereas enrichment based on cTI disease score reduced this to 234 (61% reduction), cortical thickness to 312 (47% reduction), log NfL to 303 (49% reduction), and age to 375 (37% reduction). For *GRN*, the unenriched design required 457 participants per arm, and cTI, cortical thickness, log NfL, and age enrichment reduced this to 116 (75% reduction), 124 (73% reduction), 139 (69% reduction), and 289 (37% reduction), respectively. Sample size could not be estimated for *MAPT* enrichment because too few individuals met the biomarker thresholds to support a reliable analysis

Table 2 reports the enrichment thresholds and cohort composition for each genetic group and biomarker, including the number and proportion of asymptomatic carriers who met each threshold, the number of symptomatic carriers, and the total proportion of carriers included in the hybrid enriched cohorts. Table 3 reports the analytic per-arm sample size estimates for enriched and unenriched trial designs, including values before and after accounting for 10% dropout and the percentage reduction in recruited N per arm relative to the unenriched “all carriers” scenario. For comparison, we also report the required sample size when enrichment is based solely on age, providing a simple baseline against which to evaluate biomarker-based strategies. Across both cohorts, cTI-based enrichment consistently produced the largest reductions in required sample size, followed by cortical thickness and log NfL, indicating that the cTI disease score is the most efficient enrichment metric in this setting. These reconstructed trial efficiencies were robustly confirmed by Monte Carlo simulation-based power analyses, which demonstrated that cTI-informed enrichment achieves the target 80% empirical power at substantially lower recruited sample sizes than alternative strategies for both *C9orf72* and *GRN* mutation groups (Figure 6; see Table S2 for complete simulation grid results).

**Table 2.** Enrichment thresholds and cohort composition. For each genetic group and enrichment metric, the table reports: the total number of asymptomatic carriers, the number and proportion of asymptomatic carriers who meet the enrichment threshold, the number of symptomatic carriers (always included), and the overall number and proportion of mutation carriers included in the hybrid enriched cohort. Thresholds were defined as the 80th percentile of each metric in the healthy control group.

| Genetic Group | Enrichment metric | Asymptomatic Total N | Asymptomatic Enriched N (% retained) | Symptomatic N (always included) | Total included N (% of all carriers) |
| --- | --- | --- | --- | --- | --- |
| C9orf72 | cTI Disease Score > 0.371 | 93 | 18 (19) | 36 | 54 (42) |
|  | Cortical Thickness < 2.366 | 93 | 34 (37) | 36 | 70 (54) |
|  | log NfL > 2.417 | 93 | 27 (29) | 36 | 63 (49) |
|  | Age >45 | 93 | 45 (48) | 36 | 81 (63) |
| GRN | cTI Disease Score > 0.371 | 101 | 18 (18) | 22 | 40 (33) |
|  | Cortical Thickness < 2.366 | 101 | 21(21) | 22 | 43 (35) |
|  | log NfL > 2.417 | 101 | 24 (24) | 22 | 46 (37) |
|  | Age > 45 | 101 | 60 (59) | 22 | 82 (67) |
| MAPT | cTI Disease Score > 0.371 | 44 | 15 (34) | 12 | 27 (48) |
|  | Cortical Thickness < 2.366 | 44 | 12 (27) | 12 | 24 (43) |
|  | log NfL > 2.417 | 44 | 4 (9) | 12 | 16 (29) |
|  | Age > 45 | 44 | 13 (29) | 12 | 25 (45) |

**Table 3.**
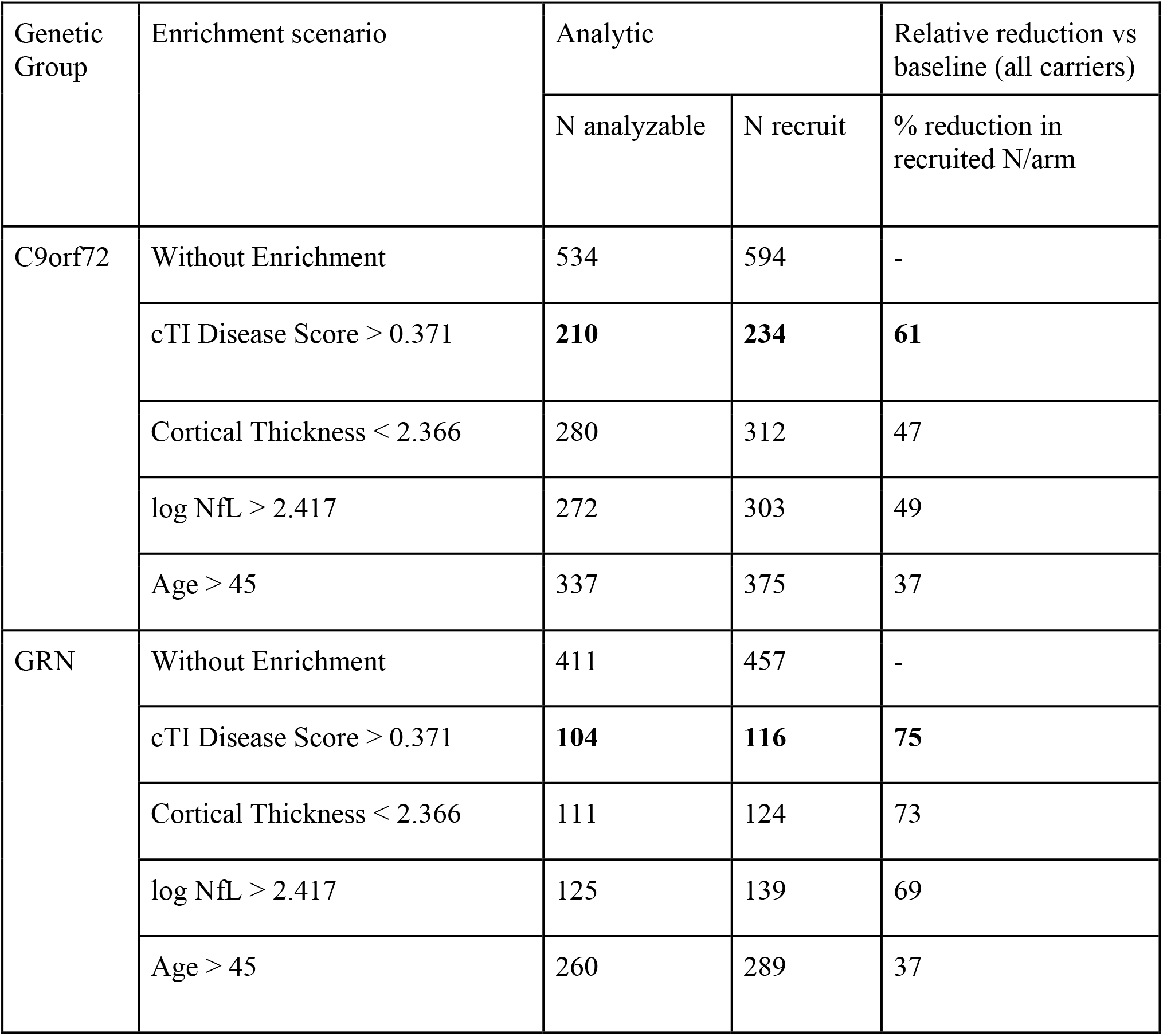
Analytic sample size estimates for enriched and unenriched trial designs. For each genetic group and recruitment strategy (no enrichment vs enrichment based on cTI disease score, cortical thickness, log NfL or age), the table reports the analytic per-arm sample size needed to detect a 50% reduction in 2-year FTLD-CDR-SOB change with 80% power at a two-sided α = 0.05. For each strategy, we show the required number of analyzable participants per arm N analyzable) and the number to recruit per arm after inflating for 10% dropout (N recruit). The percentage reduction in N recruit relative to the unenriched “all carriers” design is also provided to quantify the efficiency gains from biomarker-based enrichment.

| Genetic Group | Enrichment scenario | Analytic |  | Relative reduction vs baseline (all carriers) |
| --- | --- | --- | --- | --- |
|  |  | N analyzable | N recruit | % reduction in recruited N/arm |
| C9orf72 | Without Enrichment | 534 | 594 | - |
|  | cTI Disease Score > 0.371 | **210** | **234** | **61** |
|  | Cortical Thickness < 2.366 | 280 | 312 | 47 |
|  | log NfL > 2.417 | 272 | 303 | 49 |
|  | Age > 45 | 337 | 375 | 37 |
| GRN | Without Enrichment | 411 | 457 | - |
|  | cTI Disease Score > 0.371 | **104** | **116** | **75** |
|  | Cortical Thickness < 2.366 | 111 | 124 | 73 |
|  | log NfL > 2.417 | 125 | 139 | 69 |
|  | Age > 45 | 260 | 289 | 37 |

**Figure 6.**
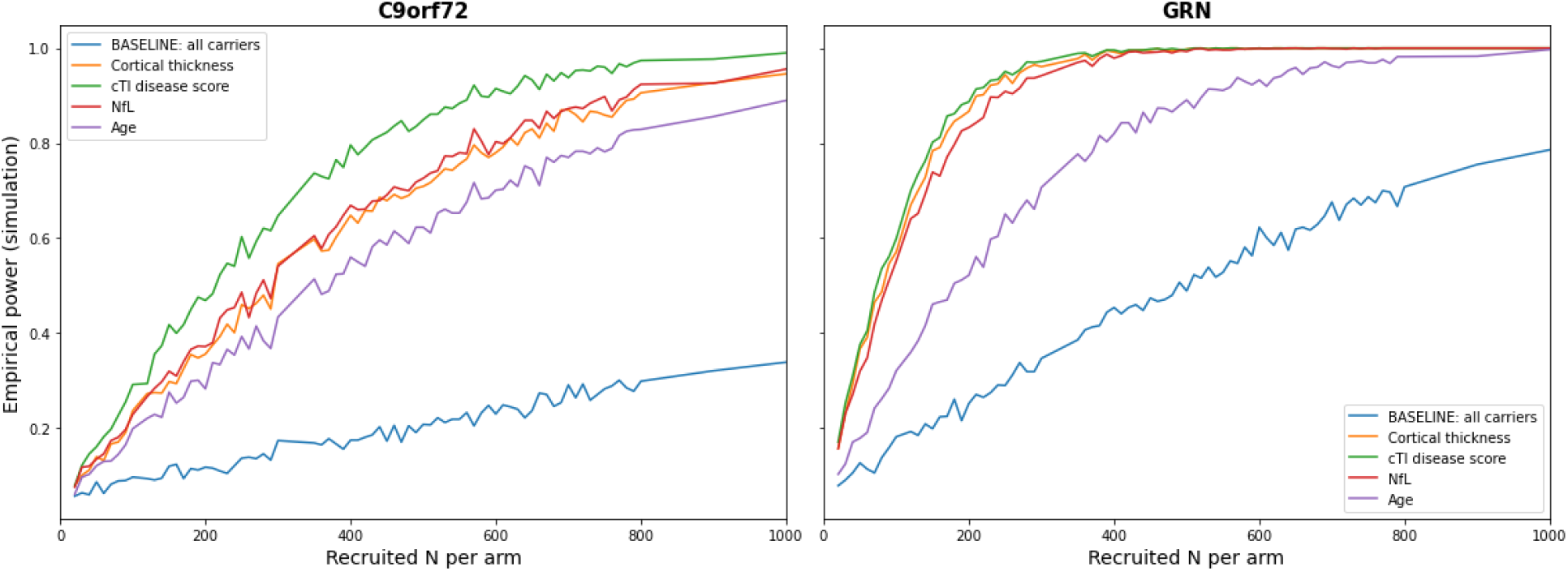
Power versus recruited sample size for enriched and unenriched designs. Each panel shows Monte Carlo–estimated power as a function of recruited sample size per arm for one genetic group (*C9orf72, GRN*). Curves compare an unenriched design (recruiting all mutation carriers) with designs enriched using cTI disease score, cortical thickness, log-transformed plasma NfL, or age. Across both groups, cTI-based enrichment achieves 80% power at lower sample sizes than the unenriched design, with cortical thickness and NfL providing intermediate gains.

## 4 DISCUSSIONS

In this study, we developed and validated a personalized disease stratification framework for genetic FTD that accounts for substantial heterogeneity in disease progression. Using contrastive trajectory inference (cTI) applied to structural MRI data from the GENFI cohort, we identified disease-specific patterns of brain atrophy, assigned individualized disease scores to mutation carriers, and discovered distinct disease subtypes characterized by markedly different progression rates. While previous studies have applied data-driven approaches to FTD progression modeling, our work uniquely combines subtype discovery with individualized progression modeling and demonstrates practical utility for clinical trial enrichment. Critically, we showed that cTI-derived disease scores substantially improve clinical trial enrichment strategies for the asymptomatic phase, requiring fewer total participants to achieve adequate statistical power compared to conventional biomarkers including cortical thickness and neurofilament light chain.

Traditional disease scoring approaches in neurodegenerative disorders typically assume a single, uniform progression trajectory, which fails to capture the heterogeneity observed in diseases like FTD where multiple distinct progression pathways coexist [6]. In contrast, cTI explicitly models disease heterogeneity by identifying multiple trajectories within the same dataset through machine learning. By doing so, it enables subtype-aware scoring that more accurately locates each individual along their disease course and supports stratified analyses in clinical trials that may reveal treatment effects obscured by population averaging. This conceptual shift from a single trajectory to multiple data-driven trajectories provides a mechanistic explanation for why cTI-based enrichment can outperform single-biomarker approaches in terms of required sample size.

These findings align with previous observations of phenotypic heterogeneity in FTD and extend prior work by providing quantitative, biomarker-based evidence for biologically distinct disease variants. Unlike earlier classification schemes based primarily on clinical phenotypes, our data-driven approach identifies subtypes based on patterns of neuroanatomical change, potentially capturing underlying pathophysiological differences that may require different therapeutic approaches. [27]

The cTI-derived disease scores showed robust construct validity through strong and significant correlations with established clinical and cognitive measures, including the Boston Naming Test, Digit Symbol Substitution Test, verbal fluency, MINI-SEA total score, Trail Making Test Part B, FTD-CDR Sum of Boxes, Mini-Mental State Examination, and estimated years to onset. These correlations reflected clinically meaningful relationships, as indicated by the scores’ ability to track longitudinal clinical progression.

Our subtyping analysis revealed that macrostructural brain changes in FTD mutation carriers follow two distinct biological trajectories. By applying contrastive trajectory inference, we separated what has traditionally been viewed as a single asymptomatic phase into a Progressive Track (Subtype 2) and a Stable Track (Subtype 3). Despite only a small difference in age (Figure S1), there is a dissociation between asymptomatic individuals that are on a progressing track (Subtype 2), from asymptomatic individuals (Subtype 3) that have brain changes not related to active axonal injury (neurofilament light chain elevation) and clinical-cognitive decline. The dissociation between the imaging-derived disease score and the clinical state in Subtype 3 may represent a resilient state. In these individuals, the brain structural variations detected by T1-weighted imaging likely represent benign neuroanatomical variation or developmental/aging differences rather than the onset of the FTD pathological cascade. Further, some symptomatic individuals were classified in Subtype 3, suggesting that they might be slowly progressive cases in the future.

Hierarchical regression analysis confirmed that baseline subtype membership significantly predicts future decline in processing speed and language, independent of baseline clinical severity, highlighting the prognostic relevance of these imaging-derived tracks. This has immediate implications for clinical trial design. Subtype 3 carriers, despite having pathogenic mutations, appear biologically stable over the periods typically studied in trials. Including these stable carriers in disease-modifying therapy trials could substantially dilute the observed treatment effect, as they do not yet exhibit the active neurodegeneration (high neurofilament light chain) that these therapies are intended to arrest.

A key translational result of this study is that cTI disease scores could substantially improve the efficiency of clinical trial recruitment in genetic FTD when including asymptomatic subjects. Our power analyses, based on assumptions of a 50% treatment effect on FTD-CDR Sum of Boxes over a two-year period with 80% power and a 5% significance level, showed that cTI-based enrichment required fewer participants than enrichment based on cortical thickness or neurofilament light chain for both *C9orf72* and *GRN* mutation carriers. This result is consistent with emerging evidence from Alzheimer disease trials, where multivariate staging biomarkers have also improved trial efficiency compared with single-modality markers.[28]. While less asymptomatic subjects met the set entry criteria for cTI than for NfL or cortical thickness, cTI allowed for better patient selection compensated by a smaller requirement for sample size.

This improvement appears to be driven by several features of the cTI framework. First, cTI scores capture patterns of covariation across multiple brain regions that univariate biomarkers cannot, providing a far more comprehensive assessment of disease stage than focal atrophy measures. This capability is directly illustrated by the top feature contributors driving the pseudotemporal progression axis, which span distinct structural pathologies (Table S1).

Specifically, the model relies on macrostructural subcortical changes (such as inferior lateral ventricles volume) alongside prominent cortical markers (mean cortical thickness) and frontal white matter hyperintensities. By simultaneously capturing these complementary metrics, the framework yields a highly robust, representative index of overall disease progression. Second, cTI-based enrichment selects asymptomatic individuals with higher disease scores, indicating greater proximity to clinical conversion, which increases the average rate of clinical decline in the enrolled cohort and reduces outcome variance. Third, cTI scores are explicitly optimized to maximize disease-specific variance while removing normal aging effects, making them particularly sensitive to pathological change.

In rare diseases such as FTD, where patient populations are geographically dispersed and recruitment is logistically challenging, even modest reductions in required sample size can determine trial feasibility[8,29]. Recruitment difficulties commonly lead to long recruitment periods and delayed trial completion, which inevitably increases overall trial costs [7,29]. Moreover, enrolling asymptomatic individuals at high risk of near-term conversion enables prevention trials, which may be more effective than symptomatic interventions if neurodegeneration becomes irreversible after symptom onset [5]. cTI-based enrichment therefore directly addresses one of the major barriers to therapeutic development in FTD by identifying asymptomatic cases at high risk of progression, along with the creation of a unified disease score allowing efficient merging of symptomatic and asymptomatic cases in the same clinical trial.

Several methodological strengths support the validity of our findings. We used rigorous image preprocessing and data harmonization to reduce scanner-related variance while preserving biological variability. Careful data cleaning excluded potentially confounding cases, covariate adjustment controlled for demographic factors, and semi-supervised learning minimized bias from predefined trajectory assumptions. However, several limitations should be considered. Although GENFI is the largest genetic FTD cohort worldwide, sample sizes for individual mutation groups remain modest, particularly for *MAPT* carriers. While we validated disease scores using longitudinal clinical data, longitudinal MRI data were insufficient to directly model neuroimaging trajectories over time, so we relied on cross-sectional trajectory inference. Our trial simulations also assumed uniform treatment effects across disease scores, although real-world efficacy may vary by disease score or subtype.

Future work should address these limitations and extend our findings. First, investigating the biological mechanisms underlying the two disease subtypes could provide mechanistic insight and identify subtype-specific therapeutic targets, including possible differences in proteinopathy, network dysfunction, neuroinflammatory profiles, and genetic modifiers. Second, incorporating additional imaging modalities (functional MRI, diffusion MRI, PET) and fluid biomarkers into the cTI framework may further improve disease scoring accuracy and subtype separation. Third, prospective clinical trials are needed to empirically test the predicted sample size reductions and evaluate the practical benefits of cTI-based enrichment strategies. Finally, extending the subtyping and scoring algorithm to make fuller use of longitudinal data could improve trajectory modeling and allow more precise prediction of individual disease progression. Prospective longitudinal validation in independent cohorts would strengthen confidence in subtype stability and score trajectories.

In conclusion, we developed a personalized disease progression scoring framework for genetic FTD that captures disease heterogeneity using machine learning applied to structural MRI data. cTI-derived disease scores that could constitute a robust clinical trial enrichment strategy requiring fewer participants to achieve adequate statistical power than conventional biomarkers. By modeling disease heterogeneity rather than averaging over it, personalized scoring approaches may help accelerate progress toward effective treatments for FTD and related neurodegenerative disorders.

## 5 Acknowledgements

We would like to thank all participants and their families for taking part in the GENFI study.

## 6 Funding

This study was supported by multiple funding sources. S.D. receives salary funding from the Fond de Recherche du Québec - Santé (FRQS). GENFI2 is funded by the Canadian Institutes for Health Research. J.B.R. is supported by the Medical Research Council (MC_UU_00030/14; MR/T033371/1), Wellcome Trust (220258), and the National Institute for Health and Care Research Cambridge Biomedical Research Centre (NIHR203312: the views expressed are those of the authors and not necessarily those of the National Institute for Health and Care Research or the Department of Health and Social Care). M.M., E.F., S.D., and R.L. Jr. have received funding from two Canadian Institutes of Health Research project grants (MOP-327387 and PJT-175242) and from the Weston Brain Institute for the conduct of this study.

## 7 Competing interests

The authors report no competing interests.

## 9 Appendix Coinvestigators

### GENFI Consortium Members

Annabel Nelson, Martina Bocchetta, David Cash, David L Thomas, Emily Todd, Hanya Benotmane, Jennifer Nicholas, Kiran Samra, Rachelle Shafei, Carolyn Timberlake, Thomas Cope, Timothy Rittman, Antonella Alberici, Enrico Premi, Roberto Gasparotti, Valentina Cantoni, Emanuele Buratti, Andrea Arighi, Chiara Fenoglio, Elio Scarpini, Giorgio Fumagalli, Vittoria Borracci, Giacomina Rossi, Giorgio Giaccone, Giuseppe Di Fede, Paola Caroppo, Pietro Tiraboschi, Sara Prioni, Veronica Redaelli, David Tang-Wai, Ekaterina Rogaeva, Miguel Castelo-Branco, Morris Freedman, Ron Keren, Sandra Black, Sara Mitchell, Christen Shoesmith, Robart Bartha, Rosa Rademakers, Janne M. Papma, Lucia Giannini, Rick van Minkelen, Yolande Pijnenburg, Benedetta Nacmias, Camilla Ferrari, Cristina Polito, Gemma Lombardi, Valentina Bessi, Michele Veldsman, Christin Andersson, Hakan Thonberg, Linn Öijerstedt, Vesna Jelic, Paul Thompson, Tobias Langheinrich, Albert Lladó, Anna Antonell, Jaume Olives, Mircea Balasa, Nuria Bargalló, Sergi Borrego-Ecija, Ana Verdelho, Ana Gorostidi, Jorge Villanua, Marta Cañada, Mikel Tainta, Miren Zulaica, Myriam Barandiaran, Patricia Alves, Benjamin Bender, Lisa Graf, Annick Vogels, Mathieu Vandenbulcke, Philip Van Damme, Rose Bruffaerts, Koen Poesen, Pedro Rosa-Neto, Serge Gauthier, Anne Bertrand, Aurélie Funkiewiez, Daisy Rinaldi, Dario Saracino, Olivier Colliot, Sabrina Sayah, Catharina Prix, Elisabeth Wlasich, Olivia Wagemann, Sandra Loosli, Sonja Schönecker, Tobias Hoegen, Jolina Lombardi, Sarah Anderl-Straub, Adeline Rollin, Gregory Kuchcinski, Maxime Bertoux, Thibaud Lebouvier, Vincent Deramecour, Beatriz Santiago, Diana Duro, Maria João Leitão, Maria Rosario Almeida, Miguel Tábuas-Pereira, Sónia Afonso

## 11 Supplementary

### Data Cleaning and Control Group Definition

To ensure a rigorously defined healthy control group, we applied a multi-step data-cleaning procedure restricted to non-carrier participants. First, individuals with any evidence of clinical impairment were excluded: all non-carriers with an FTLD-CDR-SOB score greater than zero were removed (n = 61). Participants with missing CDR values were retained to avoid unnecessary loss of otherwise healthy controls.

Because both plasma neurofilament light chain (NfL) and total white matter hyperintensity (WMH) volume exhibited strong positive skew, outlier detection was performed using a log-transformed interquartile-range (IQR) method. Briefly, biomarker values were log-transformed, and the upper outlier threshold was defined as:

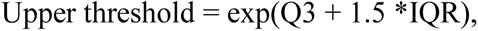

where (Q3) is the 75th percentile and IQR = (Q3 - Q1) of the log-transformed distribution. Using this approach, participants exceeding the upper thresholds for plasma NfL (25.94 pg/mL) and WMH volume (50486.17 mm³) were excluded. This resulted in the removal of 9 NfL outliers and 5 WMH outliers. All remaining non-carrier participants constituted the final healthy control sample used in downstream analyses.

### Visit Selection for Healthy Controls

Because the number of available visits varied across participants, only a single visit was retained for each healthy control to avoid biasing subsequent models toward individuals with more repeated measurements. When multiple visits were available, the visit with a valid plasma NfL measurement was selected; if several such visits existed, the most recent visit was chosen. For participants without any NfL data, the most recent available visit was retained.

### Covariate Adjustment

To remove the effects of demographic variables on imaging and biomarker measures, all features were residualized for age, sex, and education. For each feature, a linear regression model was fitted in the healthy control group using age, gender, and years of education as predictors. The resulting regression coefficients were then applied to all participants, and adjusted values were computed as the residuals (observed − predicted), yielding covariate-adjusted measurements independent of these demographic factors.

### Visit Selection for Mutation Carriers

For participants carrying pathogenic mutations, only a single visit per individual was retained to ensure consistent representation across subjects with differing follow-up durations. When only one visit was available, that visit was selected. For participants with multiple visits, we first ordered visits chronologically and excluded the final visit to ensure a subsequent follow-up existed for downstream analyses. Among the remaining visits, priority was given to those with an available FTLD-CDR-SOB score; if multiple eligible visits remained, we further prioritized visits with available plasma NfL. When ties persisted, the most recent visit among the eligible set was chosen.

### Merging Left and Right Hemisphere Features

Following covariate adjustment, left and right hemisphere features were combined to reduce dimensionality and increase statistical power, given the limited sample sizes inherent to rare disease cohorts such as genetic FTD. Although hemispheric collapsing represents a methodological limitation, it was necessary to ensure model stability while retaining biologically meaningful information. Cortical thickness measures were averaged across hemispheres for each region, whereas volumetric measures, including subcortical volumes, cortical gray matter volume, and regional WMH burden, were summed across hemispheres. In addition to these merged features, two asymmetry indices were computed to retain information about potential lateralization effects: the ratio of left-to-right WMH burden and the ratio of left-to-right mean cortical thickness. This approach reduced the total number of imaging features from 120 to 67 (44% reduction) while preserving both regional specificity and hemispheric asymmetry patterns relevant to FTD.

### Feature with highest contribution in cTI

**Table S1.** Top 10% feature contributors to the contrastive trajectory inference (cTI) score. This table lists features with the highest cTI contributions after adjustment for age, sex, and education, with inferior lateral ventricle volume and non–white-matter hypointensities showing the largest weights

| Feature | cTI contribution |
| --- | --- |
| Inferior lateral ventricles volume | 159 |
| Non–white-matter hypointensities volume | 151 |
| Frontal white-matter hyperintensities volume | 40.7 |
| Mean cortical thickness | 37.4 |
| Pallidum volume | 17.8 |
| Hippocampus volume | 16.2 |

### Comparison of Age Distributions in cTI Subtypes

**Figure S1.**
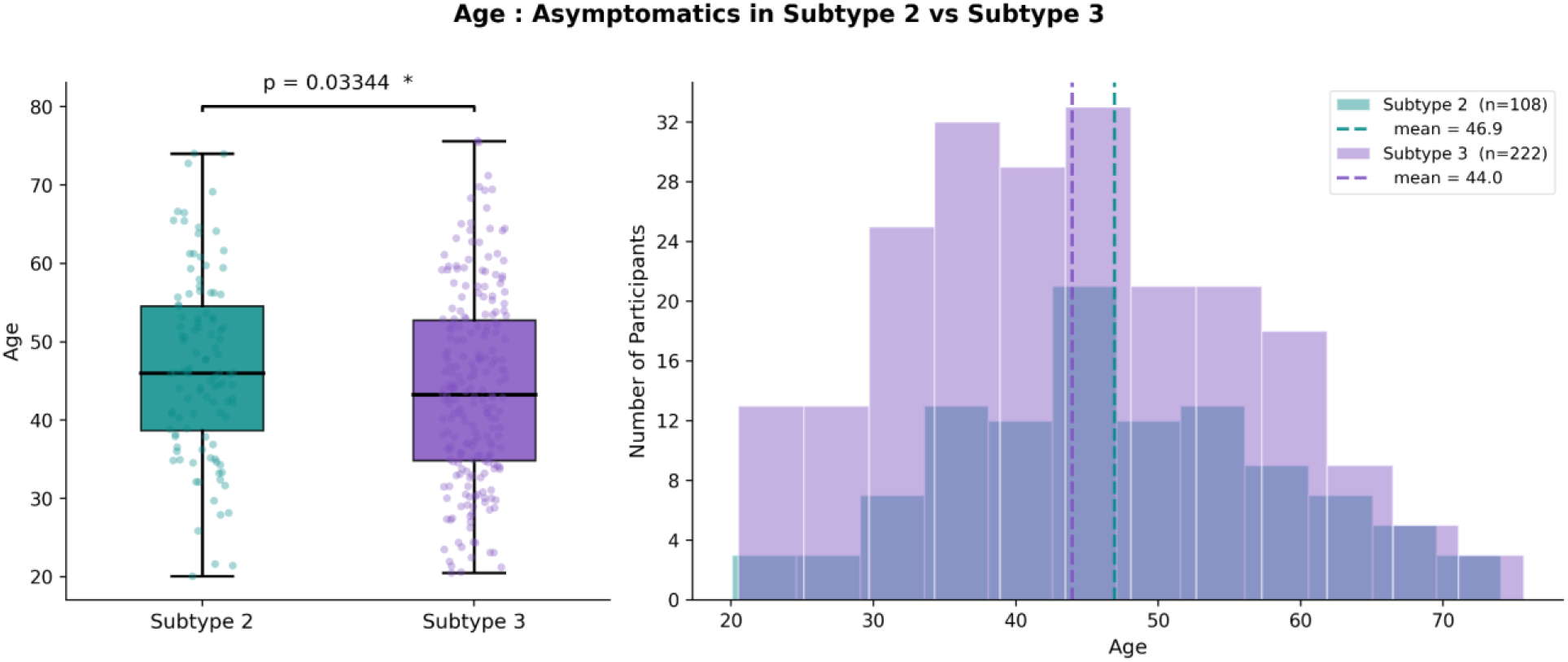
Age distribution of asymptomatic participants in subtype 2 and subtype 3. Boxplots and overlaid histograms show that asymptomatic individuals in subtype 2 are slightly older than those in subtype 3 (mean ages 46.9 vs 44.0 years), with a significant difference on the Mann–Whitney U test (p = 0.033).

### Prognostic Utility of Subtype Membership

To evaluate whether the identified neuroimaging subtypes capture clinically relevant disease heterogeneity, we tested the prognostic utility of baseline subtype membership in predicting longitudinal clinical decline. Individual rates of change (slopes) were calculated for seven cognitive and functional measures using linear regression of test scores against age across all available longitudinal visits (minimum 2 timepoints per participant).

We employed hierarchical regression models to determine if subtype classification provides incremental predictive value beyond baseline disease severity. The Baseline Model (Model 1) included only the baseline clinical score and chronological age. The Full Model (Model 2) added subtype classification as a categorical predictor. The incremental predictive value was quantified using the change in variance explained (ΔR^2^), F-tests for nested model comparison, and partial correlations (r_p_) controlling for baseline severity. To ensure the generalizability of these prognostic effects, models were evaluated using 5-fold cross-validation (ΔR^2^_CV_). All predictors were standardized prior to analysis to allow for comparison of effect sizes.

### Subtype-Specific Prediction of Longitudinal Decline

Analysis of 470 mutation carriers with longitudinal data (range: 201–321 participants per measure) revealed that subtype membership provides significant incremental predictive value for specific cognitive domains, independent of baseline clinical assessment.

#### Domain-Specific Prognostic Value

The prognostic utility of subtype classification was most pronounced in the domains of processing speed and language:

- **Processing Speed:** Subtype membership most strongly predicted decline in the Digit Symbol test (ΔR^2^ = 0.025, p = 0.004; partial r = 0.201, p < 0.001). Cross-validation confirmed robust out-of-sample prediction for this domain (ΔR^2^_CV = 0.015).
- **Language:** Significant incremental predictive value was observed for the Boston Naming Test (ΔR^2^=0.013, p=0.035, partial r=0.163, p=0.003).
- **Executive and Functional Domains:** Marginally significant associations emerged for executive function (TMT-B: ΔR^2^ = 0.011, p = 0.066) and functional impairment (FTLD-CDR-SB: ΔR^2^ = 0.010, p = 0.052). Notably, both measures showed significant partial correlations (ranging from −0.13$ to −0.14, p < 0.05) when controlling for baseline severity.

#### Global and Social Cognition

No significant incremental predictive value was observed for global cognition (MMSE), verbal fluency, or social cognition measures (Mini-SEA).

The modest but significant effect sizes (*ΔR*^2^ ≈ 1 − 2.5%) demonstrate that subtype classification captures biologically meaningful phenotypic variation that is not fully accounted for by standard clinical staging. These results suggest that accounting for subtype-specific trajectories provides complementary prognostic information, particularly for identifying individuals at risk for rapid decline in processing speed and language.

### Simulation-based power analysis for enrichment

The simulation-based power analysis used the same mutation groups, enrichment strategies, recruitment cohorts, and longitudinal dataset as in the analytic power analysis described in the main Methods (i.e., identical trial scenario, endpoint, design assumptions, and annualized FTLD-CDR-SOB change).

For a given mutation group, enrichment strategy, and candidate per-arm sample size n, we ran 1000 simulated trials using the following procedure: (1) sampled 2n participants with replacement from the cohort of interest (baseline or enriched hybrid); (2) randomized them 1:1 to active vs placebo; (3) generated placebo endpoints as baseline FTLD-CDR-SOB plus a simulated 2-year change, obtained by fitting a mean model to the resampled cohort, computing residual standard deviation, and drawing new noise from a normal distribution with that standard deviation; (4) imposed a 50% treatment effect by subtracting *δ* = 0.50 × *μ_placebo_* from the active arm endpoints, where *μ_placebo_* is the mean annualized change of the placebo bootstrap sample scaled to 2 years and varies across simulated trials; (5) applied dropout by removing each participant independently with probability equal to the assumed drop rate; and (6) compared 2-year change between arms using a two-sample t-test (two-sided α = 0.05).

Empirical power for each *n* was defined as the proportion of simulated trials with p < 0.05. We evaluated a grid of per-arm sample sizes and, for each mutation group and enrichment strategy, defined the required sample size as the smallest *n* achieving at least 80% empirical power.

**Table S2.** Simulation-based sample size estimates for enriched and unenriched trial designs. For each genetic group and enrichment scenario, the table reports the minimum recruited sample size per arm (from a predefined grid) required to achieve ≥80% power in Monte Carlo simulations, along with the empirical power at that sample size. Simulations used the same trial assumptions as the analytic power analysis and accounted for observed variability in 2-year FTLD-CDR-SOB change and dropout.

| Genetic Group | Enrichment scenario | Simulation |  |
| --- | --- | --- | --- |
|  |  | n_min_per_arm (recruited) | achieved power at n_min |
| C9orf72 | Without Enrichment | 3100 | 80.5 |
|  | cTI Disease Score > 0.371 | **430** | 80.7 |
|  | Cortical Thickness < 2.366 | 620 | 81.1 |
|  | log NfL > 2.417 | 570 | 83 |
|  | Age > 45 | 770 | 81.6 |
| GRN | Without Enrichment | 1100 | 83 |
|  | cTI Disease Score > 0.371 | **150** | 80.2 |
|  | Cortical Thickness < 2.366 | 170 | 82.4 |
|  | log NfL > 2.417 | 190 | 82.6 |
|  | Age > 45 | 380 | 81.6 |

## Notes

### Competing Interest Statement

The authors have declared no competing interest.

## REFERENCES

1. Greaves CV, Rohrer JD. An update on genetic frontotemporal dementia. J Neurol. 2019 Aug;266(8):2075–86. doi:10.1007/s00415-019-09363-4

2. Antonioni A, Raho EM, Lopriore P, Pace AP, Latino RR, Assogna M, et al. Frontotemporal Dementia, Where Do We Stand? A Narrative Review. Int J Mol Sci. 2023 Jul 21;24(14):11732. doi:10.3390/ijms241411732 PubMed PMID: 37511491; PubMed Central PMCID: PMC10380352.

3. Häkkinen S, Chu SA, Lee SE. Neuroimaging in genetic frontotemporal dementia and amyotrophic lateral sclerosis. Neurobiology of Disease. 2020 Nov 1;145:105063. doi:10.1016/j.nbd.2020.105063

4. Young AL, Marinescu RV, Oxtoby NP, Bocchetta M, Yong K, Firth NC, et al. Uncovering the heterogeneity and temporal complexity of neurodegenerative diseases with Subtype and Stage Inference. Nat Commun. 2018 Oct 15;9(1):1. doi:10.1038/s41467-018-05892-0

5. Rohrer JD, Nicholas JM, Cash DM, van Swieten J, Dopper E, Jiskoot L, et al. Presymptomatic cognitive and neuroanatomical changes in genetic frontotemporal dementia in the Genetic Frontotemporal dementia Initiative (GENFI) study: a cross-sectional analysis. The Lancet Neurology. 2015 Mar 1;14(3):253–62. doi:10.1016/S1474-4422(14)70324-2

6. Staffaroni AM, Quintana M, Wendelberger B, Heuer HW, Russell LL, Cobigo Y, et al. Temporal order of clinical and biomarker changes in familial frontotemporal dementia. Nat Med. 2022 Oct;28(10):2194–206. doi:10.1038/s41591-022-01942-9 PubMed PMID: 36138153; PubMed Central PMCID: PMC9951811.

7. Bartoshyk P, O’Caoimh R. Update on Disease-Modifying Pharmacological Treatments for Frontotemporal Dementia (FTD): A Scoping Review of Registered Trials. NeuroSci. 2025 Dec;6(4):114. doi:10.3390/neurosci6040114

8. Boxer AL, Gold M, Feldman H, Boeve BF, Dickinson SLJ, Fillit H, et al. New directions in clinical trials for frontotemporal lobar degeneration: Methods and outcome measures. Alzheimer’s & Dementia. 2020;16(1):131–43. doi:10.1016/j.jalz.2019.06.4956

9. Tsai RM, Boxer AL. Therapy and clinical trials in frontotemporal dementia: past, present, and future. Journal of Neurochemistry. 2016;138(S1):211–21. doi:10.1111/jnc.13640

10. Staffaroni AM, Ljubenkov PA, Kornak J, Cobigo Y, Datta S, Marx G, et al. Longitudinal multimodal imaging and clinical endpoints for frontotemporal dementia clinical trials. Brain. 2019 Feb;142(2):443–59. doi:10.1093/brain/awy319 PubMed PMID: 30698757; PubMed Central PMCID: PMC6351779.

11. Zammitt D, Brotherhood EV, Fearn C, Greaves C, Hayes O, Harding E, et al. Barriers and Facilitators to Participation in Clinical Trials Related to Familial Frontotemporal Dementia: A Qualitative Study. Molecular Genetics & Genomic Medicine. 2024;12(11):e70038. doi:10.1002/mgg3.70038

12. Bocchetta M, Todd EG, Bouzigues A, Cash DM, Nicholas JM, Convery RS, et al. Structural MRI predicts clinical progression in presymptomatic genetic frontotemporal dementia: findings from the GENetic Frontotemporal dementia Initiative cohort. Brain Commun. 2023 Apr 1;5(2):fcad061. doi:10.1093/braincomms/fcad061

13. Soltaninejad M, Iturria-Medina Y, Rajabli R, Bezgin G, Hosseini-Kamkar N, Bouzigues A, et al. MRI-based classifier to identify close-to-onset cases in C9orf72 genetic frontotemporal dementia [Internet]. bioRxiv; 2025 [cited 2026 Mar 8]. p. 2025.10.18.683237. Available from: https://www.biorxiv.org/content/10.1101/2025.10.18.683237v1 doi:10.1101/2025.10.18.683237

14. Soltaninejad M, Dadar M, Collins DL, Rajabli R, Venkatraghavan V, Bouzigues A, et al. White matter hyperintensities precede other biomarkers in GRN frontotemporal dementia. Alzheimer’s & Dementia. 2025;21(10):e70695. doi:10.1002/alz.70695

15. Dutt S, Leichter D, Cobigo Y, Wolf A, Kornak J, Clark A, et al. Individualized Atrophy-Based Prediction of Dementia Progression in Familial Frontotemporal Lobar Degeneration With Bayesian Linear Mixed-Effects Modeling. Annals of Neurology. n/a(n/a). doi:10.1002/ana.78167

16. Planche V, Mansencal B, Manjon JV, Tourdias T, Catheline G, Coupé P, et al. Anatomical MRI staging of frontotemporal dementia variants. Alzheimer’s & Dementia. 2023;19(8):3283–94. doi:10.1002/alz.12975

17. McCarthy J, Borroni B, Sanchez-Valle R, Moreno F, Laforce R, Graff C, et al. Data-driven staging of genetic frontotemporal dementia using multi-modal MRI. Human Brain Mapping. 2022 Apr 15;43(6):1821–35. doi:10.1002/hbm.25727

18. Pengo M, Premi E, Borroni B. Dissecting the Many Faces of Frontotemporal Dementia: An Imaging Perspective. IJMS. 2022 Oct 25;23(21):12867. doi:10.3390/ijms232112867

19. Iturria-Medina Y, Khan AF, Adewale Q, Shirazi AH, the Alzheimer’s Disease Neuroimaging Initiative. Blood and brain gene expression trajectories mirror neuropathology and clinical deterioration in neurodegeneration. Brain. 2020 Feb 1;143(2):661–73. doi:10.1093/brain/awz400

20. Iturria-Medina Y, Carbonell F, Assadi A, Adewale Q, Khan AF, Baumeister TR, et al. Integrating molecular, histopathological, neuroimaging and clinical neuroscience data with NeuroPM-box. Commun Biol. 2021 May 21;4(1):1. doi:10.1038/s42003-021-02133-x

21. Wilke C, Reich S, van Swieten JC, Borroni B, Sanchez-Valle R, Moreno F, et al. Stratifying the Presymptomatic Phase of Genetic Frontotemporal Dementia by Serum NfL and pNfH: A Longitudinal Multicentre Study. Annals of Neurology. 2022;91(1):33–47. doi:10.1002/ana.26265

22. Dadar M, Collins DL. BISON: Brain tissue segmentation pipeline using T1 -weighted magnetic resonance images and a random forest classifier. Magn Reson Med. 2021 Apr;85(4):1881–94. doi:10.1002/mrm.28547 PubMed PMID: 33040404.

23. Johnson WE, Li C, Rabinovic A. Adjusting batch effects in microarray expression data using empirical Bayes methods. Biostatistics. 2007 Jan 1;8(1):118–27. doi:10.1093/biostatistics/kxj037

24. Abid A, Zhang MJ, Bagaria VK, Zou J. Exploring patterns enriched in a dataset with contrastive principal component analysis. Nat Commun. 2018 May 30;9(1):2134. doi:10.1038/s41467-018-04608-8

25. Gordon E, Rohrer JD, Kim LG, Omar R, Rossor MN, Fox NC, et al. Measuring disease progression in frontotemporal lobar degeneration. Neurology. 2010 Feb 23;74(8):666–73. doi:10.1212/WNL.0b013e3181d1a879 PubMed PMID: 20177120; PubMed Central PMCID: PMC2830919.

26. Morris JC, Weintraub S, Chui HC, Cummings J, Decarli C, Ferris S, et al. The Uniform Data Set (UDS): clinical and cognitive variables and descriptive data from Alzheimer Disease Centers. Alzheimer Dis Assoc Disord. 2006;20(4):210–6. doi:10.1097/01.wad.0000213865.09806.92 PubMed PMID: 17132964.

27. Tartaglia MC, Mackenzie IRA. Recent Advances in Frontotemporal Dementia. Canadian Journal of Neurological Sciences. 2023 Jul;50(4):485–94. doi:10.1017/cjn.2022.69

28. Holland D, McEvoy LK, Desikan RS, Dale AM. Enrichment and Stratification for Predementia Alzheimer Disease Clinical Trials. PLoS One. 2012 Oct 17;7(10):e47739. doi:10.1371/journal.pone.0047739 PubMed PMID: 23082203; PubMed Central PMCID: PMC3474753.

29. Franzen S, Nuytemans K, Bourdage R, Caramelli P, Ellajosyula R, Finger E, et al. Gaps in clinical research in frontotemporal dementia: A call for diversity and disparities focused research. Alzheimers Dement. 2023 Dec;19(12):5817–36. doi:10.1002/alz.13129 PubMed PMID: 37270665; PubMed Central PMCID: PMC10693651.

